# Low hatching success in the critically endangered kākāpō (Strigops habroptilus) is driven by early embryo mortality not infertility

**DOI:** 10.1101/2020.09.14.295949

**Authors:** James L. Savage, Jodie M. S. Crane, Kākāpō Recovery Team, Nicola Hemmings

## Abstract

In many endangered species, reproductive failure is a major barrier to recovery. The critically endangered kākāpō (*Strigops habroptilus*) exemplifies this challenge: 61% of their eggs fail to hatch, and of these 73% show no sign of development. Undeveloped eggs have previously been attributed to male infertility, but recent studies of non-threatened bird species suggest fertilisation failure is rare in the wild. The underlying causes of fertilisation failure and embryo death differ, so distinguishing between them is essential for effective conservation management. Here we show that the majority (72%, n=124) of undeveloped kākāpō eggs are fertilised, and combine this with conservation programme data on natural copulations, artificial inseminations, and paternity of developed eggs, to generate the most precise estimate to date of fertility in a wild population. We also demonstrate, for the first time in a wild bird, that artificial insemination results in greater numbers of sperm reaching the egg.

## Introduction

Hatching failure is a common problem for endangered birds, with dramatic consequences for population viability (Jamieson & Ryan 2000; Heber & Briskie 2010). Hatching failure can result from either fertilisation failure or embryo mortality, but these have different proximate causes: for example, fertilisation failure may result from poor sperm quality or abnormal mating behaviour (Brillard 1990; Lifjeld et al. 2007), while embryo mortality is more likely driven by parental genetic incompatibilities/inbreeding (Sittmann et al. 1966; Hemmings et al. 2012b), female condition (Coleman & Siegel 1966; Lerner et al. 1993), and/or environmental factors (Romanoff 1949; Beissinger et al. 2005). Distinguishing fertilisation failure from early embryo mortality is therefore crucial for developing effective conservation interventions, but relatively few studies or conservation programmes have attempted to do this; most assume that undeveloped eggs are unfertilised (Jamieson & Ryan 2000; Hemmings et al. 2012a).

Distinguishing embryo mortality from infertility is especially challenging if mortality occurs in the first few days of development, which in well-studied domestic poultry is when most embryos die (Christensen 2001). At this stage the embryo is a tiny disc of cells, easily mistaken for an unfertilised blastodisc on visual inspection (Kosin 1944), and hence traditional macroscopic methods of examining eggs can over-estimate fertilisation failure in undeveloped eggs. The few studies that have microscopically inspected unhatched eggs suggest that early embryo death may be the main cause of hatching failure in wild birds (e.g. Hemmings and Evans 2020), but such studies remain uncommon, particularly in threatened populations where hatching failure is of greatest concern.

The kākāpō (*Strigops habroptilus*) is a large, flightless, nocturnal parrot endemic to New Zealand. Once common throughout New Zealand, kākāpō rapidly declined after human settlement due to introduced predators, hunting, and land clearance (Powlesland et al. 2006), particularly after European colonisation (Bergner et al. 2016). Conservation efforts began as early as 1894, but by 1995, their total population had fallen to 51 individuals. Under intensive management kākāpō numbers have since increased, but their irregular breeding seasons and lek mating system, exacerbated by high rates of reproductive failure, pose substantial challenges for management (Elliott et al. 2006; Powlesland et al. 2006).

Kākāpō eggs often fail to hatch. Approximately 61% of kākāpō eggs have failed in the last 4 decades (401/662, 1981-2019; KRT unpublished data), much higher than the average 10-15% across birds (Koenig 1982; Spottiswoode & Møller 2004). Around 73% of failed eggs show no signs of development on visual inspection (293/401), and have previously been assumed to be unfertilised. However, without distinguishing between fertilisation failure and early embryo death, the mechanistic basis of hatching failure in kākāpō remains unclear.

This distinction would facilitate the identification of individuals or combinations of parents susceptible to infertility and/or early offspring mortality, informing management interventions such as artificial insemination and translocation.

In 2019, the largest kākāpō breeding season on record allowed us to assess true fertility rates in kākāpō for the first time, using data from two separate breeding islands and multiple clutches per female. We examined the fertility status of undeveloped eggs using methods developed previously for other birds (Hemmings et al. 2012a), and combined this with data on kākāpō copulation events in the wild, artificial inseminations, and known parentage of developed embryos to infer patterns of individual fertility and embryo survival across the entire species.

## Materials and Methods

### Study system

The entire kākāpō species (August 2020: 209 individuals) is managed on predator-free offshore islands, the breeding population (110 reproductively mature adults) primarily on Whenua Hou/Codfish Island (lat. −46.77, lon. 167.63) and Anchor Island (lat. −45.76, lon. 166.51). Kākāpō are lek breeders, exhibiting female-only parental care (Eason et al. 2006) and skewed reproduction among males (Merton et al. 1984). All mating events are recorded by individual transmitters (Wildtech New Zealand Ltd) fitted prior to the breeding season. Male transmitters continuously monitor male activity, and activate a receiver to detect nearby female transmitters whenever the preceding ten minutes of activity is ≥96% of the maximum possible activity (i.e. when males are running, fighting, or mating). While active, the receiver also records female transmitter activity, allowing non-mating encounters (with low female activity) to be excluded. Every kākāpō nest is monitored: eggs are replaced with decoys and artificially incubated, and chicks returned to nests after hatching (Elliott et al. 2001).

Kākāpō reproduce only when natural food is abundant (Powlesland & Lloyd 1994), typically every 2-4 years. Successful breeding depends on the availability of masting tree fruit, particularly podocarps such as rimu (*Dacrydium cupressinum*) (Powlesland et al. 1992). Masting in 2018/19 was exceptional, with the highest abundance of rimu fruit since records began in 1997. The 2018/19 breeding season was both the largest and earliest on record: 252 eggs were laid, the earliest on 25 December 2018, several weeks earlier than normal. The Kākāpō Recovery Team also induced many females to produce a second clutch by removing and artificially incubating/head-rearing their first clutch. Twelve females were artificially inseminated during the season.

### Egg data collection

During artificial incubation, eggs were candled daily by experienced Kākāpō Recovery Team members and removed from the incubator after 5 days if no embryo was visible. We opened undeveloped eggs, removed the contents, then separated the yolks and preserved them in 5% formalin solution, following Hemmings et al. (2012a). In consultation with Ngāi Tahu (New Zealand South Island Māori iwi), we then shipped the preserved yolks for analysis at the University of Sheffield, UK (CITES institution GB041). In total, we obtained samples from 128 undeveloped eggs: 98% of all undeveloped eggs laid in 2019. Eggs that failed later in development (i.e. with a visible embryo) were not used in this study, but embryos were staged and DNA-sampled to ascertain paternity as part of the Kākāpō Recovery Team’s standard protocols.

### Egg fertility and sperm numbers

We removed yolks from formalin and rinsed them with phosphate buffered saline solution (PBS). We located the germinal disc on the yolk, removed the overlaying perivitelline layer (PVL) using dissecting scissors, and washed it in PBS before mounting on a microscope slide. We also tried to remove the germinal disc, but in many cases it remained strongly adhered to the PVL so we examined them together. Some yolks disintegrated prior to fixation; in these cases we examined pieces of PVL found within the sample.

We examined the PVL for sperm and germinal disc samples for embryonic cells by staining them with the fluorescent DNA marker Hoechst 33342 (Thermo Fisher Scientific), and inspecting them under a fluorescence microscope with a BP 340-380 excitation filter and LP 425 suppression filter (Birkhead et al. 2008). The number of sperm reaching the avian ovum is related to (a) the number inseminated (Wishart 1987) and (b) the likelihood of hatching (Eslick & McDaniel 1992; Hemmings & Birkhead 2015), so we also counted the number of sperm on known areas of PVL (±1mm^2^). We classified eggs as either fertilised (embryonic nuclei and sperm present) or unfertilised (no embryonic nuclei and few/no sperm). In total, we established the fertility status of 124/128 (97%) undeveloped eggs; four eggs were too deteriorated to reliably examine.

### Statistical methods

Using generalised linear mixed models, we analysed the effects of clutch (first or second), female multiple mating (one or multiple males), and artificial insemination (yes/no) on (a) egg fertility and (b) the number of PVL sperm. We specified a binomial distribution and logit link function in the analysis of egg fertility as the response variable was binary (fertile, yes/no), and a Poisson distribution and log link in the analysis of sperm numbers as the response variable was a count (offset = log(PVL area)). We included female identity as a random effect in both models, and compared model fits using AIC and Likelihood Ratio Tests. Data from third clutches were excluded because only a single female produced a third clutch. Further methodological details and analysis code are available in supplementary material.

## Results

### Fertilisation failure and embryo mortality in undeveloped eggs

Of 57 reproductively mature female kākāpō, 49 nested in 2019 on Anchor Island and Whenua Hou / Codfish Island. Nests of 36 females were removed to induce a second clutch, and of these 30 re-nested. Across all clutches, 119/252 eggs were visibly fertile. The rest (133) remained undeveloped, and would previously have been assumed to be unfertilised. A total of 86 eggs hatched and 72 chicks reached juvenile age (150 days) (**Figure 1**).

**Figure 1:**
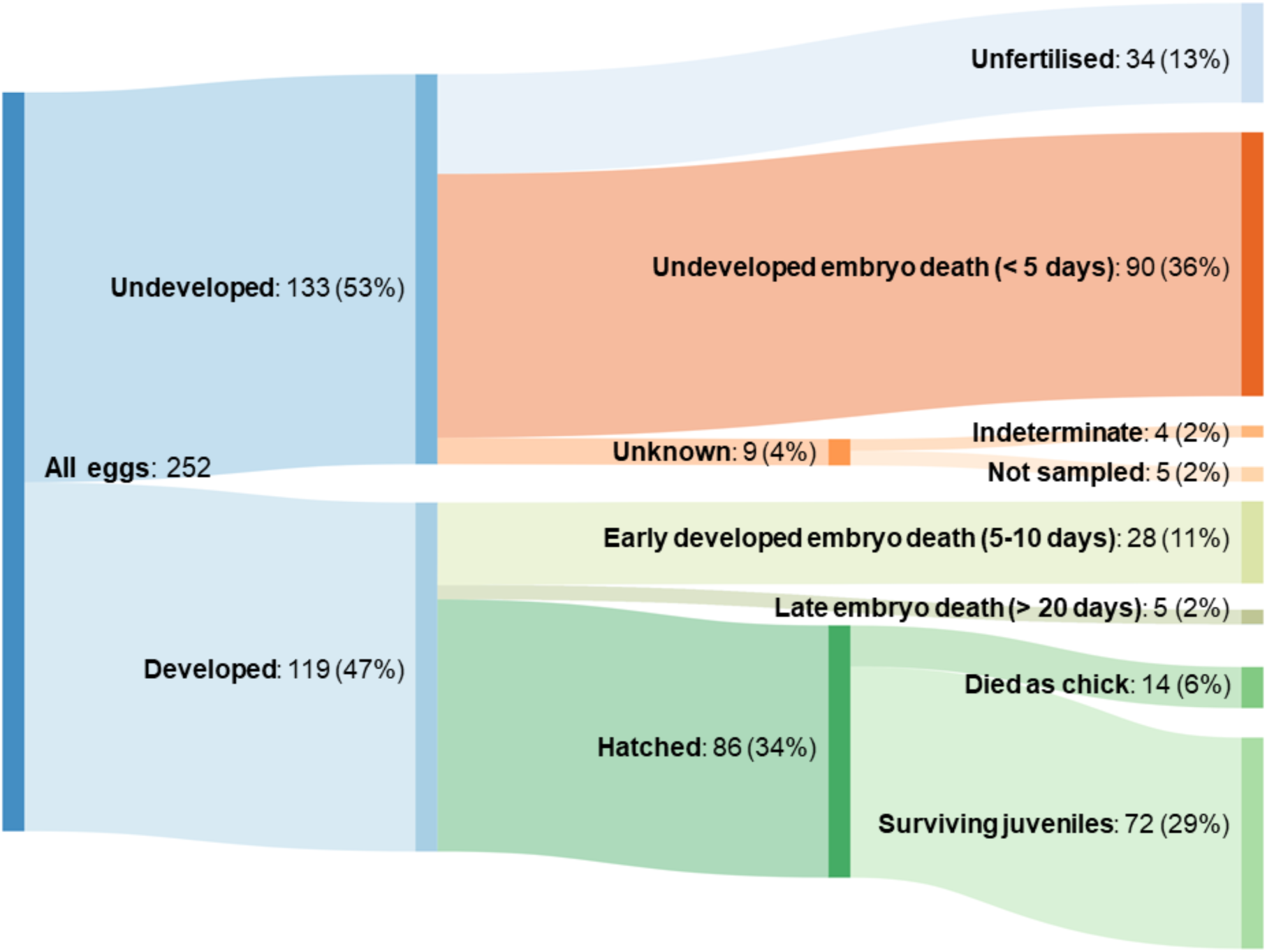
Visualisation of fates of all 252 eggs laid during the 2019 breeding season. Approximately equal numbers of eggs were laid on Anchor Island (123 from 21 females) and Whenua Hou / Codfish Island (129 from 28 females), and there were no differences in egg fertility or embryo death between eggs from different islands. No embryos died during the middle third of the incubation period (10-20 days). Some eggs laid on one island were fostered to the other island following artificial incubation.

Of 128 undeveloped eggs examined, we found that 90 were fertilised and 34 were unfertilised. Excluding the four eggs that could not be examined due to degradation, 73% (90/124) were fertilised. The true rate of fertilisation failure across all eggs was hence 14% (34/248), considerably lower than 52% (131/252) which would have been assumed without examination of undeveloped eggs. Consequently, the primary driver of hatching failure (and overall reproductive failure) in kākāpō in 2018/19 was very early embryo mortality (< day 5 of development), which occurred in at least 36% (90/252) of all eggs laid (**Figure 1**). Almost half of all eggs (118/252; 47%) failed due to embryo death in the first third of the incubation period (days 0-10).

Fertilisation failure was more common in first than second clutches (estimate = 1.473, z = 2.427, p (|z|) = 0.015; **Figure 2**). We found no evidence that females that copulated with more than one male had greater egg fertilisation success (estimate = 0.612, z = 0.871, p (|z|) = 0.384), but artificially inseminated females may have had increased fertilisation success (estimate = 2.124, z = 1.664, p (|z|) = 0.096). The most parsimonious model included clutch and artificial insemination, but a model that also included multiple mating had a similar fit (ΔAICC 1.244, p (χ2) = 0.385). Proportions of developed eggs were similar across both islands.

**Figure 2:**
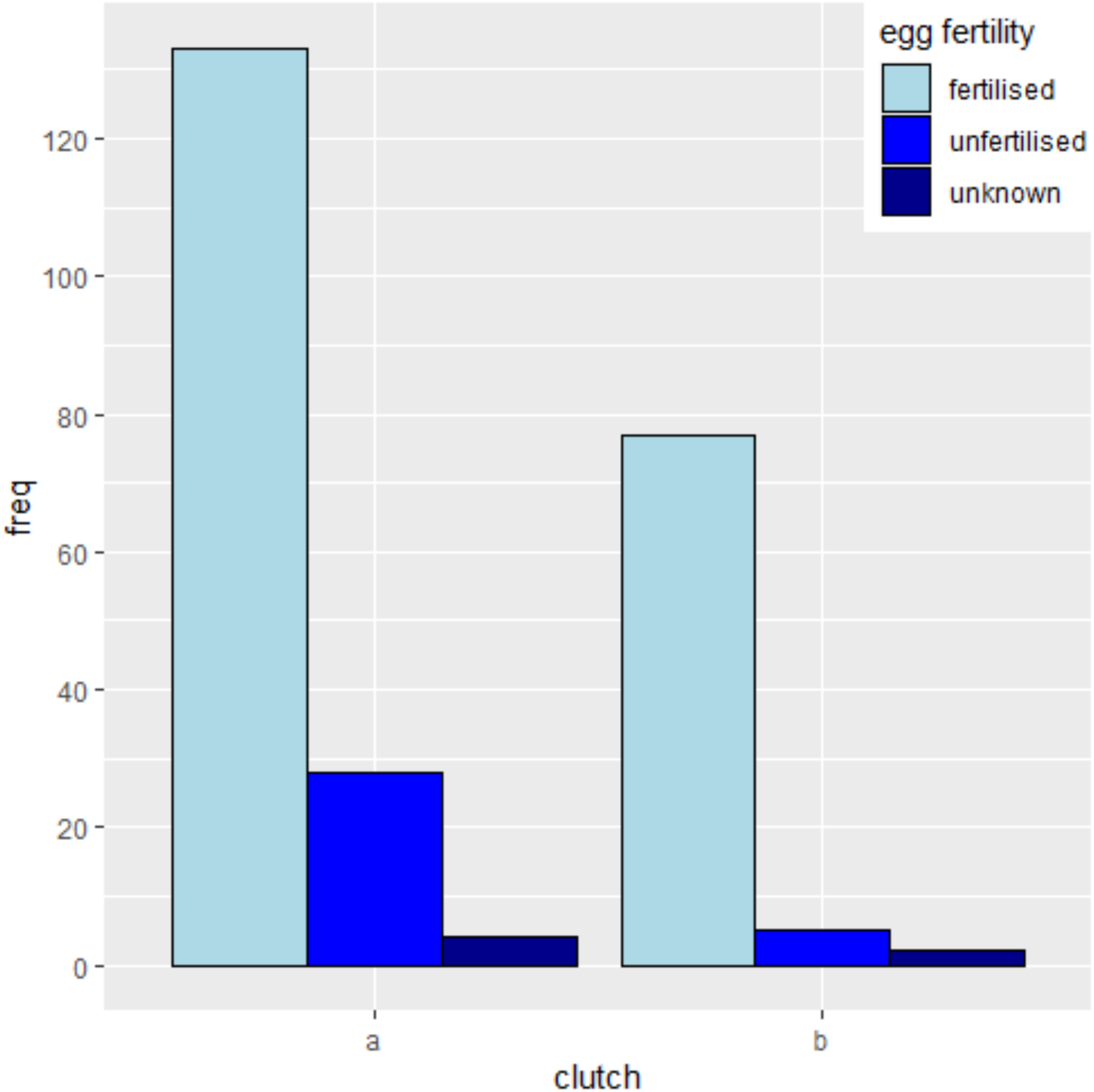
The frequency of fertilised and unfertilised eggs across first (a) and second (b) clutches. Fertilisation failure was more common in first clutches.

### Individual fertility

Of 31 males known to have copulated (or used for artificial insemination), 26 (84%) fertilised at least one egg. Most males were therefore not infertile/sterile, because at least one of their insemination attempts resulted in successful fertilisation. A total of 18 males produced embryos that reached at least day five of development (based on KRT data from developed eggs). However, five male kākāpō showed no clear evidence of fertility in 2019 despite copulating with females and/or being used for artificial insemination (4 of these males have produced chicks in previous years). Two of these males inseminated females that produced fertilised but undeveloped eggs of unknown paternity, however their mates also copulated with fertile males and in some cases produced developed eggs of known paternity with other males. It is therefore unlikely that these two males fertilised the eggs laid by these females.

Across all breeding females, only one produced no fertilised eggs in 2019. Of the other 48 females, 10 (21%) produced eggs in which all the embryos died at an early undeveloped stage, and three of these females have produced chicks in previous years. Based on these data, we estimate that individual infertility rates in reproductively active kākāpō in 2018/19 were approximately 17% in males and 2% in females.

### Sperm numbers reaching eggs

We counted the number of sperm on the PVL of 83 fertilised eggs. The number of sperm per cm^2^ of PVL ranged from 0-82 (median = 15), but these numbers are likely to be conservative because the PVL was often obscured by adhered yolk and/or embryonic tissue (as a result of the sample fixation process). We did not analyse the number of holes made by sperm entering the ovum, because the inner PVL was frequently absent or degraded. We found no evidence that PVL sperm numbers varied between first and second clutches (estimate = 0.018, z = 0.220, p (|z|) = 0.826). However, eggs from multiply mated females may have had more sperm (estimate = −0.268, z = −1.905, p (|z|) = 0.057), and artificially inseminated females exhibited significantly more sperm (estimate = −2.091, z = −5.214, p (|z|) <0.001) (**Figure 3**). The best-fitting model included artificial insemination and multiple mating, however a model including only artificial insemination was not clearly inferior (ΔAIC 1.666, p (χ2) = 0.055).

**Figure 3:**
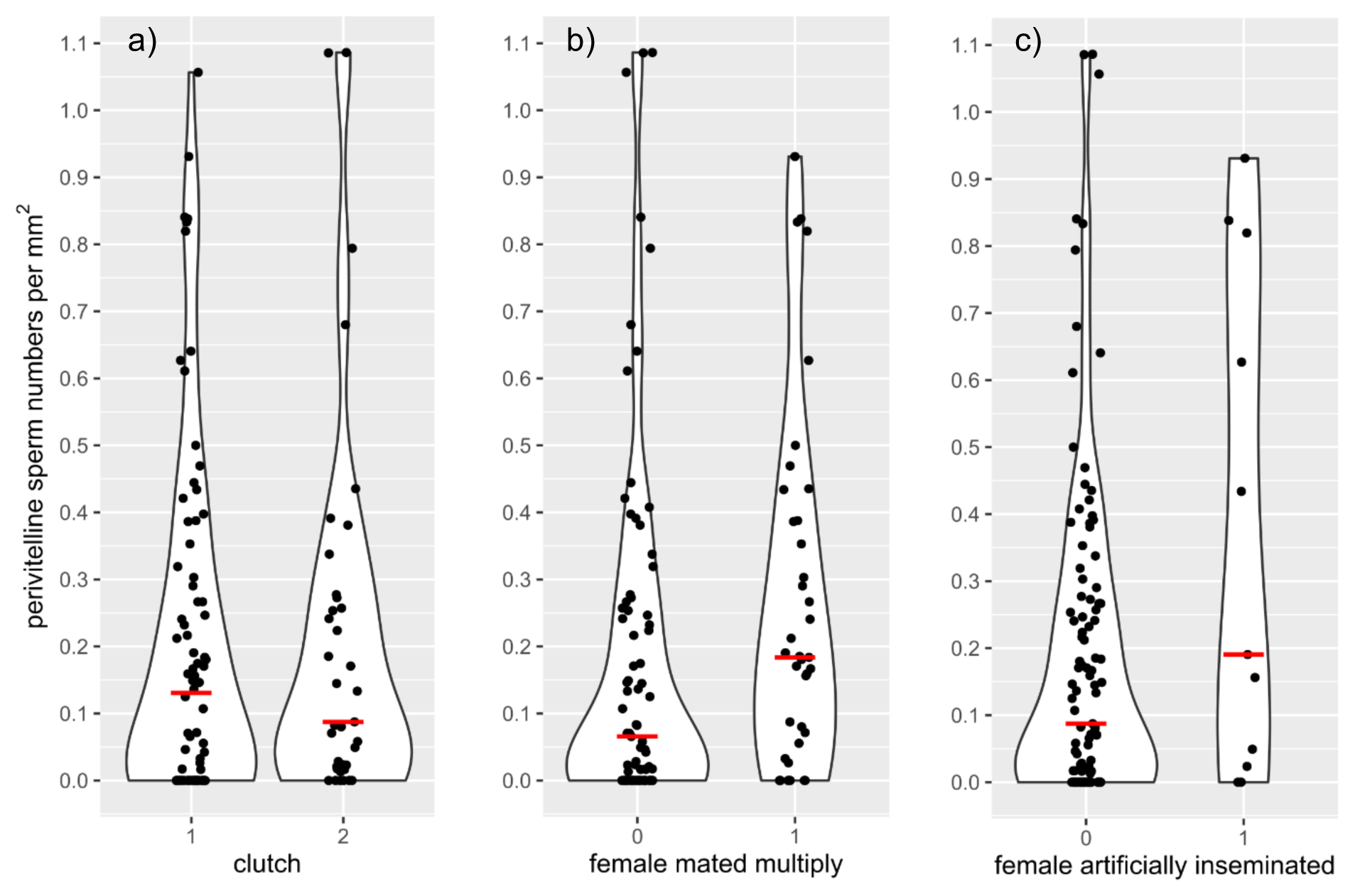
The number of sperm (per mm^2^) counted on the perivitelline layer (PVL) of undeveloped eggs, compared between (a) clutches 1 and 2, (b) females that didn’t (0) or did (1) mate with multiple males, and (c) females that weren’t (0) or were (1) artificially inseminated. Horizonal lines show median values. There were no significant differences in sperm numbers between clutches 1 and 2 (z = −0.20, p=0.84), but eggs from multiply-mated females may have had more sperm (z = 1.87, p=0.061) and eggs from artificially inseminated females had significantly more sperm (z = 5.19, p < 0.0001).

## Discussion

Kākāpō are critically endangered, and their recovery is hampered by exceptionally high rates of hatching failure and presumed infertility. Our results show that early embryo mortality is the main driver of kākāpō reproductive failure: almost three-quarters of apparently unfertilised kākāpō eggs are in fact fertilised, but die at an early stage of development. True individual infertility rates in kākāpō are hence lower than previously thought, with at least 83% of breeding males being capable of fertilisation and 98% of breeding females producing fertilised eggs. Our results also demonstrate the potential benefits of artificial insemination as a management tool in wild populations, as eggs laid by artificially inseminated females had much higher sperm numbers on the perivitelline layer.

### Embryo mortality as a driver of reproductive failure in kākāpō

Only 51 kākāpō existed in 1995, and all but one of this founder population originated from a relatively small and potentially inbred population from Rakiura / Stewart Island (Bergner et al. 2014). The risk of inbreeding after this bottleneck is exacerbated by the species’ lek breeding system, with local paternity dominated by few males. A previous study found no link between male heterozygosity and fertility in male kākāpō (White et al. 2015), however that study assumed all undeveloped eggs were unfertilised. Moreover, early embryo death is more likely to be related to heterozygosity of the embryo itself than that of either parent, and is an expected consequence of inbreeding as inbred individuals express more lethal recessives during development (Charlesworth & Willis 2009). In contrast, the evidence that inbreeding depresses adult sperm function and individual fertility in wild populations is equivocal (Losdat et al. 2014, 2018). It therefore seems likely that early embryo mortality in kākāpō is largely driven by inbreeding depression on early survival.

### Identification of infertile individuals and consequences for breeding management

In wild populations it is often impossible to identify infertile individuals, as copulations are rarely monitored and fertilisation failure is largely misidentified. Most long-term population studies, for example, only collect breeding information on individuals that provide care and/or produce offspring; identifying males who copulate but fail to father or care for offspring is practically impossible. The unique management regime of kākāpō, with copulations remotely recorded and eggs artificially incubated, hence represents a unique opportunity to accurately assess individual fertility across an entire species.

Although our results indicate that individual fertility rates in kākāpō are higher than previously assumed, we still identified five males exhibiting no evidence of fertilisation success despite copulation attempts. In absolute terms this is a relatively small number of individuals, but still amounts to 17% of breeding kākāpō males: a significant proportion of the potential genetic diversity available. Investigating whether male infertility is a result of physiological (e.g. sperm production), behavioural, and/or genetic factors will be a valuable step towards understanding if and how these individuals can contribute reproductively to the population in future.

We also identified a significant number of individuals (male and female) whose offspring never developed beyond the earliest stages of embryo development (< day 5 of development). Since early embryo death is likely to be associated with inbreeding depression, an important next step will be to explore whether parental genetic similarity is related to the likelihood of early embryo survival, to inform selection of individuals for artificial insemination and/or translocation.

We found clear evidence that more sperm reached the eggs of artificially inseminated females, and weaker evidence that a higher proportion of these females’ eggs were fertilised. This is encouraging, as only 12 females were artificially inseminated, and procedure effectiveness is likely to have improved through the breeding season. This relatively new conservation management tool could play an important role maximising future reproductive success and genetic diversity in kākāpō.

### Timing of breeding and consequences for fertility

The early 2018/9 breeding season allowed the Kākāpō Recovery Team to induce second clutches, almost doubling the reproductive capacity of the population. We found that fewer eggs were fertilised from first than from second clutches, indicating that individual fertility was lower earlier in the season; perhaps because females were unable to obtain sufficient sperm, or because males had not reached peak sperm production capacity. In many species the environmental and social cues that trigger reproduction are complex, and responses to these cues may differ between males and females (Ball & Ketterson 2008). This raises the concerning possibility that as species shift their timing of breeding in response to global warming (e.g. Hällfors *et al*. 2020), infertility might increase through sex-specific physiological constraints and resulting differences in reproductive timing.

### Conclusions

Our results demonstrate that early embryo mortality is the major driver of reproductive failure in kākāpō, with true infertility affecting a smaller but nonetheless significant proportion of individuals. Our findings both clarify the underlying reproductive problems of kākāpō, and have broader applicability to other conservation programmes. By incorporating the methods used in this study into ongoing management protocols, conservation programmes can provide a targeted individual-level response to reproductive failure. In kākāpō, individual-specific fertility and compatibility data will be crucial for selecting individuals for artificial inseminations and translocations. For species bred in captivity, this information could also inform decisions about which individuals should be paired.

Revealing this degree of individual-specific information on fertility across an entire wild population was only possible because of the extensive data on kākāpō behaviour, reproductive success, and life-history collected through their conservation management. Beyond the direct benefits to ecosystem recovery and persistence, long-term programmes such as Kākāpō Recovery generate datasets on parentage, mating behaviour, movement ecology, habitat preferences, and reproductive investment (among other topics) comparable to those of long-term research-focused studies, which are invaluable to our understanding of endangered species management and behavioural ecology more generally (Ewen et al. 2013), As long-term individual-based studies are disproportionately valuable but rare because of a dearth in long-term funding (Clutton-Brock & Sheldon 2010; Hughes et al. 2017), and conservation organisations can seldom dedicate staff to full-time research, effective collaboration between conservation professionals and researchers can greatly benefit both parties.

## Acknowledgements

Kākāpō Recovery Team staff contributing to this research: Deidre Vercoe (Operations Manager); Daryl Eason (Technical Advisor); Andrew Digby (Scientific Advisor); Karen Andrew (Supervisor); Jodie Crane (Senior Ranger); Bronwyn Jeynes (Advocacy & Logistics Ranger); Theo Thompson (Site Management & Logistics Ranger); Sara Larcombe, Freya Moore, Jake Osborne, Margie Grant, Brodie Philp, Nicki van Zyl, Liam Bolitho, Bryony Hitchcock, Leigh Joyce, Jinty MacTavish, Rachel Rouse (Field Rangers).

We thank Ngāi Tahu for their support of this project, and are grateful to the DOC permissions team and Invercargill / Murihiku office staff for administrative support.

## Author contributions

Conceptualization: JC, NH, JS; Data curation: JS, NH, KRT; Funding acquisition: JS, NH, JC; Investigation: NH, JC, JS, KRT; Methodology: NH, JS; Project administration: JS, NH; Resources: NH, KRT; Visualization: JS, NH; Writing – original draft: JS, NH; Writing – review & editing: NH, JS, JC, KRT. All authors gave final approval for publication and agree to be held accountable for the work performed therein.

## Ethics

The project was conducted as part of the New Zealand Department of Conservation Kākāpō Recovery Programme’s conservation management activities, with approval from Ngāi Tahu.

## References

Ball, G.F. & Ketterson, E.D. (2008). Sex differences in the response to environmental cues regulating seasonal reproduction in birds. Philos. Trans. R. Soc. B Biol. Sci., 363, 231–246.

Beissinger, S.R., Cook, M.I. & Arendt, W.J. (2005). The shelf life of bird eggs: Testing egg viability using a tropical climate gradient. Ecology, 86, 2164–2175.

Bergner, L.M., Dussex, N., Jamieson, I.G. & Robertson, B.C. (2016). European colonization, not polynesian arrival, impacted population size and genetic diversity in the critically endangered New Zealand Kakapo. J. Hered., 107, 593–602.

Bergner, L.M., Jamieson, I.G. & Robertson, B.C. (2014). Combining genetic data to identify relatedness among founders in a genetically depauperate parrot, the Kakapo (Strigops habroptilus). Conserv. Genet., 15, 1013–1020.

Birkhead, T.R., Hall, J., Schut, E. & Hemmings, N. (2008). Unhatched eggs: Methods for discriminating between infertility and early embryo mortality. Ibis, 150, 508–517.

Brillard, J.P. (1990). Control of fertility in birds. INRA.

Charlesworth, D. & Willis, J.H. (2009). The genetics of inbreeding depression. Nat. Rev. Genet., 10, 783–796.

Christensen, V.L. (2001). Factors associated with early embryonic mortality. Worlds. Poult. Sci. J., 57, 359–372.

Clutton-Brock, T. & Sheldon, B.C. (2010). Individuals and populations: The role of long-term, individual-based studies of animals in ecology and evolutionary biology. Trends Ecol. Evol., 25, 562–573.

Coleman, J.W. & Siegel, P.B. (1966). Selection for Body Weight at Eight Weeks of Age: 5. Embryonic Stage at Oviposition and Its Relationship to Hatchability. Poult. Sci., 45, 1008–1011.

Eason, D.K., Elliott, G.P. & Merton, D. V. (2006). Breeding biology of kakapo (Strigops habroptilus) on offshore island sanctuaries, 1990-2002. Notornis, 53, 27–36.

Elliott, G.P., Eason, D.K., Jansen, P.W., Merton, D. V, Harper, G.A. & Moorhouse, R.J. (2006). Productivity of kakapo (Strigops habroptilus) on offshore island refuges. Notornis, 53, 138–142.

Elliott, G.P., Merton, D. V & Jansen, P.W. (2001). Intensive management of a critically endangered species: The kakapo. Biol. Conserv., 99, 121–133.

Eslick, M.L. & McDaniel, G.R. (1992). Interrelationships between fertility and hatchability of eggs from broiler breeder hens. J. Appl. Poult. Res., 1, 156–159.

Ewen, J.G., Adams, L. & Renwick, R. (2013). New Zealand Species Recovery Groups and their role in evidence-based conservation. J. Appl. Ecol., 50, 281–285.

Hällfors, M.H., Antão, L.H., Itter, M., Lehikoinen, A., Lindholm, T., Roslin, T. & Saastamoinen, M. (2020). Shifts in timing and duration of breeding for 73 boreal bird species over four decades. PNAS, 117, 18557–18565.

Heber, S. & Briskie, J. V. (2010). Population bottlenecks and increased hatching failure in endangered birds. Conserv. Biol., 24, 1674–1678.

Hemmings, N. & Birkhead, T.R. (2015). Polyspermy in birds?: sperm numbers and embryo survival. Proc. R. Soc. B, 282, 20151682.

Hemmings, N. & Evans, S. (2020). Unhatched eggs represent the invisible fraction in two wild bird populations. Biol. Lett., 16, 20190763.

Hemmings, N., West, M. & Birkhead, T.R. (2012a). Causes of hatching failure in endangered birds. Biol. Lett., 8, 964–7.

Hemmings, N.L., Slate, J. & Birkhead, T.R. (2012b). Inbreeding causes early death in a passerine bird. Nat. Commun., 3, 863.

Hughes, B.B., Beas-Luna, R., Barner, A.K., Brewitt, K., Brumbaugh, D.R., Cerny-Chipman, E.B., Close, S.L., Coblentz, K.E., De Nesnera, K.L., Drobnitch, S.T., Figurski, J.D., Focht, B., Friedman, M., Freiwald, J., Heady, K.K., Heady, W.N., Hettinger, A., Johnson, A., Karr, K.A., Mahoney, B., Moritsch, M.M., Osterback, A.M.K., Reimer, J., Robinson, J., Rohrer, T., Rose, J.M., Sabal, M., Segui, L.M., Shen, C., Sullivan, J., Zuercher, R., Raimondi, P.T., Menge, B.A., Grorud-Colvert, K., Novak, M. & Carr, M.H. (2017). Long-Term studies contribute disproportionately to ecology and policy. Bioscience, 67, 271–278.

Jamieson, I.G. & Ryan, C.J. (2000). Increased egg infertility associated with translocating inbred takahe (Porphyrio hochstetteri) to island refuges in New Zealand. Biol. Conserv., 94, 107–114.

Koenig, W.D. (1982). Ecological and social factors affecting hatchability of eggs. Auk, 99, 526–536.

Kosin, I.L. (1944). Macro-and Microscopic Methods of Detecting Fertility in Unincubated Hen’s Eggs. Poult. Sci., 23, 266–269.

Lerner, S.P., French, N., McIntyre, D. & Baxter-Jones, C. (1993). Age-Related Changes in Egg Production, Fertility, Embryonic Mortality, and Hatchability in Commercial Turkey Flocks. Poult. Sci., 72, 1025–1039.

Lifjeld, J.T., Laskemoen, T., Fossøy, F., Johnsen, A. & Kleven, O. (2007). Functional infertility among territorial males in two passerine species, the willow warbler Phylloscopus trochilus and the bluethroat Luscinia svecica. J. Avian Biol., 38, 267–272.

Losdat, S., Chang, S.M. & Reid, J.M. (2014). Inbreeding depression in male gametic performance. J. Evol. Biol., 27, 992–1011.

Losdat, S., Germain, R.R., Nietlisbach, P., Arcese, P. & Reid, J.M. (2018). No evidence of inbreeding depression in sperm performance traits in wild song sparrows. Ecol. Evol., 1842–1852.

Merton, D. V, Morris, R.B. & Atkinson, I.A.E. (1984). Lek behaviour in a parrot: the kākāpō Strigops habroptilus of New Zealand. Ibis, 126, 277–283.

Powlesland, R.G. & Lloyd, B.D. (1994). Use of supplementary feeding to induce breeding in free-living kakapo Strigops habroptilus in New Zealand. Biol. Conserv., 69, 97–106.

Powlesland, R.G., Lloyd, B.D., Best, H.A. & Merton, D. V. (1992). Breeding biology of the Kakapo Strigops habroptilus on Stewart Island, New Zealand. Ibis, 134, 361–373.

Powlesland, R.G., Merton, D. V & Cockrem, J.F. (2006). A parrot apart: The natural history of the kakapo (Strigops habroptilus), and the context of its conservation management. Notornis, 53, 3–26.

Romanoff, A.L. (1949). Critical Periods and Causes of Death in Avian Embryonic Development. Auk, 66, 264–270.

Sittmann, K., Abplanalp, H. & Fraser, R.A. (1966). Inbreeding Depression in Japanese Quail. Genetics, 54, 371–379.

Spottiswoode, C. & Møller, A.P. (2004). Genetic similarity and hatching success in birds. Proc. R. Soc. B, 271, 267–272.

White, K.L., Eason, D.K., Jamieson, I.G. & Robertson, B.C. (2015). Evidence of inbreeding depression in the critically endangered parrot, the kakapo. Anim. Conserv., 18, 341–347.

Wishart, G.J. (1987). Regulation of the length of the fertile period in the domestic fowl by numbers of oviducal spermatozoa, as reflected by those trapped in laid eggs. J. Reprod. Fertil., 80, 493–498.

