## Supplementary Material for "Low hatching success in the critically endangered kākāpō (Strigops habroptilus) is driven by early embryo mortality not infertility"

#### Study System & 2018/19 Breeding Season

##### *Breeding Phenology & Productivity*

Kākāpō (*Strigops habroptilus*) reproduce only when natural food is abundant, on average every 2-4 years, and in typical breeding years fewer than 50% of mature females may lay eggs. To gain condition and rear chicks, kākāpō depend on (and are limited by) the fruit of masting tree species: particularly rimu (*Dacrydium cupressinum)* but also southern beech (*Lophozonia menziesii* and *Fuscospora spp.*), yellow silver pine (*Lepidothamnus intermedius*) and pink pine (*Halocarpus biformis*). The rimu fruit count in Autumn 2018 was 47% on Whenua Hou and 30% on Anchor, the highest since 1996, and 2018/19 also saw the largest beech mast in 29 years and a pink pine mast on Anchor Island. Higher-than-average mean air temperatures in December 2018 further contributed to the early breeding dates. Mean first mating dates were earlier on Anchor Island (2018-12-28) than on Whenua Hou (2019-01-10) for first clutches, but were the same for second clutches (2019-02-12). In total, 49 female kākāpō nested in the 2018/19 season (out of 50 mature females on Anchor Island and Whenua Hou), producing 252 eggs across 81 nests, with clutch sizes of 2-5.

##### *Artificial Incubation*

Artificial incubation of eggs took place in facilities on Anchor Island and Whenua Hou, primarily using Brinsea Maxi II Advance EX Incubators. These incubators have a capacity of 14, but to minimise risks from incubator failure many incubators were used with small numbers of kākāpō eggs in each.

#### Analysis & Visualization

Analyses were performed using generalised linear mixed models (package = lme4, function = glmer) in R version 3.6.0 (Bates et al. 2015; R Core Team 2019). Figure 1 was made using SankeyMATIC; Figures 2 and 3 were made using the R package ‘ggplot2’ (Wickham 2016), and utility functions from the ‘plyr’ package (Hadley Wickham 2011). Code for this analysis is reproduced below.

##### *R Code for Analyses*

###### Load data & Packages

```
library(lme4) # required package

setwd(choose.dir()) # choose working directory containing data files

d1 <- read.csv(file.choose()) # select .csv dataset 1 (all 2018/19 breeding season kākāpō eggs)

d2 <- read.csv(file.choose()) # select .csv dataset 2 (undeveloped 2018/19 kākāpō eggs
      examined at University of Sheffield)
```

###### Fertility Status of Eggs

```
m1a <- glmer(fertile ~ clutch + female_multiple_males + female_AI + (1|mother), family =
binomial, data = d1[d1$clutch!="c",]) # Full model; Third clutches removed as only a
single female laid a third clutch

summary(m1a) # Full model summary

m1b <- glmer(fertile ~ clutch + female_AI + (1|mother), family = binomial, data =
d1[d1$clutch!="c",]) # Clutch & AI model

m1c <- glmer(fertile ~ clutch + (1|mother), family = binomial, data = d1[d1$clutch!
="c",]) # Clutch only model

AIC(m1a,m1b,m1c) # Compare model AICs

anova(m1a,m1b) # Full comparison between two best models
```

###### Sperm Numbers on the PVL

```
m2a <- glmer(PVL_sperm ~ clutch + female_multiple_males + female_AI +
offset(log(PVL_area_mm)) + (1|mother), family = poisson, data = d2[d2$clutch!="c",]) #
Full model

summary(m2a) # full model summary

m2b <- glmer(PVL_sperm ~ + female_multiple_males + female_AI + offset(log(PVL_area_mm)) +
```

```
(1|mother), family = poisson, data = d2[d2$clutch!="c",]) # Multiple males & AI model

m2c <- glmer(PVL_sperm ~ + female_AI + offset(log(PVL_area_mm)) + (1|mother), family =
poisson, data = d2[d2$clutch!="c",]) # AI only model

AIC(m2a,m2b,m2c) # Compare model AICs

anova(m2b,m2c) # Full comparison between two best models
```

#### *R Code for Figures*

##### Figure 2

```
library(plyr); library(ggplot2) # required packages
fertbyclutch <- count(d1[d1$clutch!="c",],c("fertile","clutch"))
fertbyclutch$fertile <- mapvalues(fertbyclutch$fertile, from = c("0", "1", NA), to =
    c("unfertilised", "fertilised", "unknown")) # recode data
ggplot(fertbyclutch, aes(x=clutch, y=freq, fill=fertile)) +
     geom_bar(stat="identity", position=position_dodge(), colour = "black") +
     scale_y_continuous(breaks=seq(0,140,20)) +
     scale_fill_manual(values = c("light blue", "blue", "dark blue") ) +
     labs(fill = "egg fertility") +
     theme(legend.position = c(0.89, 0.89))
```

##### Figure 3a

```
ggplot(d2[d2$clutch!="c",], aes(x=clutch, y=mean_sperm_mm)) +
      labs(y= bquote("perivitelline sperm numbers per"*" "*mm^2), x = "clutch") +
      geom_violin(trim=TRUE) +
      geom_jitter(shape=16, position=position_jitter(0.1)) +
      stat_summary(fun.y=median, geom="point", size=10, shape = 95, color="red") +
      scale_y_continuous(breaks=seq(0,1.1,0.1))
```

##### Figure 3b

```
ggplot(d2[d2$clutch!="c",], aes(x=as.factor(female_multiple_males), y=mean_sperm_mm)) +
      labs(y= " ", x = "female mated multiply") +
      geom_violin(trim=TRUE) +
      geom_jitter(shape=16, position=position_jitter(0.1)) +
      stat_summary(fun.y=median, geom="point", size=10, shape = 95, color="red") +
      scale_y_continuous(breaks=seq(0,1.1,0.1))
```

##### Figure 3c

```
ggplot(d2[d2$clutch!="c",], aes(x=as.factor(female_AI), y=mean_sperm_mm)) +
      labs(y= " ", x = "female artificially inseminated") +
      geom_violin(trim=TRUE) +
      geom_jitter(shape=16, position=position_jitter(0.1)) +
      stat_summary(fun.y=median, geom="point", size=10, shape = 95, color="red") +
      scale_y_continuous(breaks=seq(0,1.1,0.1))
```

### References

Bates, D., Maechler, M., Bolker, B. & Walker, S. (2015). Package lme4. *J. Stat. Softw.*, 67, 1–91.

Hadley Wickham. (2011). The Split-Apply-Combine Strategy for Data Analysis. *J. Stat. Softw.*, 40, 1–29.

R Core Team. (2019). R: A Language and Environment for Statistical Computing.

Wickham, H. (2016). *ggplot2: Elegant Graphics for Data Analysis*. Springer-Verlag New York.
